# Widespread dysregulation of the circadian clock in human cancer

**DOI:** 10.1101/130765

**Authors:** Jarrod Shilts, Guanhua Chen, Jacob J. Hughey

**Affiliations:** Department of Biomedical Informatics, Vanderbilt University School of Medicine, Nashville, TN, United States; Department of Biostatistics, Vanderbilt University School of Medicine, Nashville, TN, United States

## Abstract

The mammalian circadian clock is a critical regulator of metabolism and cell division. Although multiple lines of evidence indicate that systemic disruption of the circadian clock can promote cancer, whether the clock is disrupted in primary human tumors is unknown. Here we used transcriptome data from mice to define a signature of the mammalian circadian clock based on the co-expression of 12 genes that form the core clock or are directly controlled by the clock. Our approach can be applied to samples that are not labeled with time of day and were not acquired over the entire circadian (24-h) cycle. We validated the clock signature in transcriptome data from healthy human tissues, then developed a metric we call the delta clock correlation distance (ΔCCD) to describe the extent to which the signature is perturbed in samples from one condition relative to another. We calculated the ΔCCD comparing human tumor and non-tumor samples from The Cancer Genome Atlas and eight independent datasets, discovering widespread dysregulation of clock gene co-expression in tumor samples. Subsequent analysis of data from clock gene knockouts in mice suggested that clock dysregulation in human cancer is not caused solely by loss of activity of clock genes. Furthermore, by analyzing a large set of genes previously inferred to be rhythmic in healthy human lung, we found that dysregulation of the clock in human lung cancer is accompanied by dysregulation of broader patterns of circadian co-expression. Our findings suggest that clock dysregulation is a common means by which human cancers achieve unrestrained growth and division, and that restoring clock function could be a viable therapeutic strategy in multiple cancer types. In addition, our approach opens the door to using publicly available transcriptome data to quantify clock disruption in a multitude of human phenotypes. Our method is available as a web application at https://hugheylab.shinyapps.io/deltaccd.

## Introduction

Daily rhythms in mammalian physiology are guided by a system of oscillators called the circadian clock *(1)*. The core clock consists of feedback loops between several genes and proteins, and based on work in mice, is active in nearly every tissue in the body *(2, 3)*. The clock aligns itself to environmental cues, particularly cycles of light-dark and food intake *(4–6)*. In turn, the clock regulates various aspects of metabolism *(7–9)* and is tightly linked to the cell cycle *(10–15)*.

Consistent with the tight connections between the circadian clock, metabolism, and the cell cycle, multiple studies have found that systemic disruption of the circadian system can promote cancer. In humans, long-term rotating shift work and night shift work, which perturb sleep-wake and circadian rhythms, have been associated with breast, colon, and lung cancer *(16–19)*. In mice, environmental disruption of the circadian system (e.g., through severe and chronic jet lag) increases the risk of breast cancer and hepatocellular carcinoma *(20, 21)*. Furthermore, both environmental and genetic disruption of the circadian system promote tumor growth and decrease survival in a mouse model of human lung adenocarcinoma *(22)*. While these studies support the link from the clock to cancer, other studies have established a link in the other direction, namely that multiple components of a tumor, including the RAS and MYC oncogenes, can induce dysregulation of the circadian clock *(23–25)*. Despite this progress, however, whether the clock is actually disrupted in human tumors has remained unclear. Given recent findings that stimulating the circadian clock slows tumor growth in a mouse model of melanoma *(26)*, it is important to determine the extent of clock disruption across human cancers, in order to delineate the general potential of anti-tumor strategies based on restoring or improving clock function.

When the mammalian circadian clock is functioning normally, clock genes and clock-controlled genes show characteristic rhythms in expression throughout the body and in vitro *(2, 3, 27)*. Measurements of these rhythms through time-course experiments have revealed that the clock is altered or perturbed in some human breast cancer cell lines *(28, 29)*. Existing computational methods for this type of analysis require that samples be labeled with time of day (or time since start of experiment) and acquired throughout the 24-h cycle *(30–32)*. Unfortunately, existing data from resected human tumors meet neither of these criteria. In this scenario, one approach might be to look for associations between the expression levels of clock genes and other biological and clinical variables. For example, in human breast cancer, the expression levels of several clock genes have been associated with metastasis-free survival (with the direction of association depending on the gene) *(33)*. However, because a functional circadian clock is marked less by the levels of gene expression than by periodic variation in gene expression, this type of analysis cannot necessarily be used to determine whether the clock is functional.

A more sophisticated approach is to assume the presence of periodic variation and to infer a cyclical ordering of samples, using methods such as Oscope or CYCLOPS *(34, 35)*. By applying CYCLOPS to transcriptome data from hepatocellular carcinoma, Anafi et al. found evidence for weaker or disrupted rhythmicity of several clock genes (as well as genes involved in apoptosis and JAK-STAT signaling) in tumor samples compared to non-tumor samples. Importantly, CYCLOPS does not require that samples be labeled with time of day, but it does require that the samples cover the entire cycle. As a result, the authors recommend that CYCLOPS be applied to datasets from humans with at least 250 samples *(35)*. In addition, although CYCLOPS can be used to infer rhythmicity in the expression of individual genes, it is not designed to quantify the differences in circadian clock function as a whole between multiple conditions.

Rather than attempting to infer an oscillation, an alternative approach might be to take advantage of the pattern of co-expression (e.g., pairwise correlation) that results from different clock genes having rhythms with different characteristic phases. Indeed, a previous study found different levels of co-expression between a few clock genes in different subtypes and grades of human breast cancer *(33)*. Although this finding was an important first step, its generalizability has been limited because the correlations in expression were not examined for all clock genes, in other human cancer types, or in healthy tissues where the circadian clock is known to be functional. Thus, a definitive answer to whether the circadian clock is functional across the spectrum of human cancers is still lacking.

The goal of this study was to determine whether the circadian clock is functional in human cancer. Using transcriptome data from mice, we defined a robust signature of the mammalian circadian clock based on the co-expression of clock genes. We validated the signature using transcriptome data from various organs in humans, then examined the extent to which the signature was perturbed in tumor compared to non-tumor samples from The Cancer Genome Atlas (TCGA) and from multiple independent datasets. Our findings suggest that the circadian clock is dysfunctional in a wide range of human cancers.

## Results

### Consistent correlations in expression of clock genes in mice

The progression of the mammalian circadian clock is marked by characteristic rhythms in gene expression throughout the body *(3)*. We hypothesized that the relative phasing of the rhythms of different genes would give rise to a characteristic pattern of correlations between genes. Such a pattern could be used to infer the activity of the clock, even in datasets in which samples are not labeled with time of day (Fig. 1A). To test this hypothesis, we first collected eight publicly available datasets of genome-wide, circadian gene expression from various mouse organs under both constant darkness and alternating light-dark cycles *(3, 12, 36–40)* (Table S1). We focused on 12 genes that are part of the core circadian clock or are directly controlled by the clock and that exhibit strong, consistently phased rhythms in expression in multiple organs *(32)*. For the rest of the manuscript, we refer to these 12 genes as “clock genes.”

**Figure 1.**
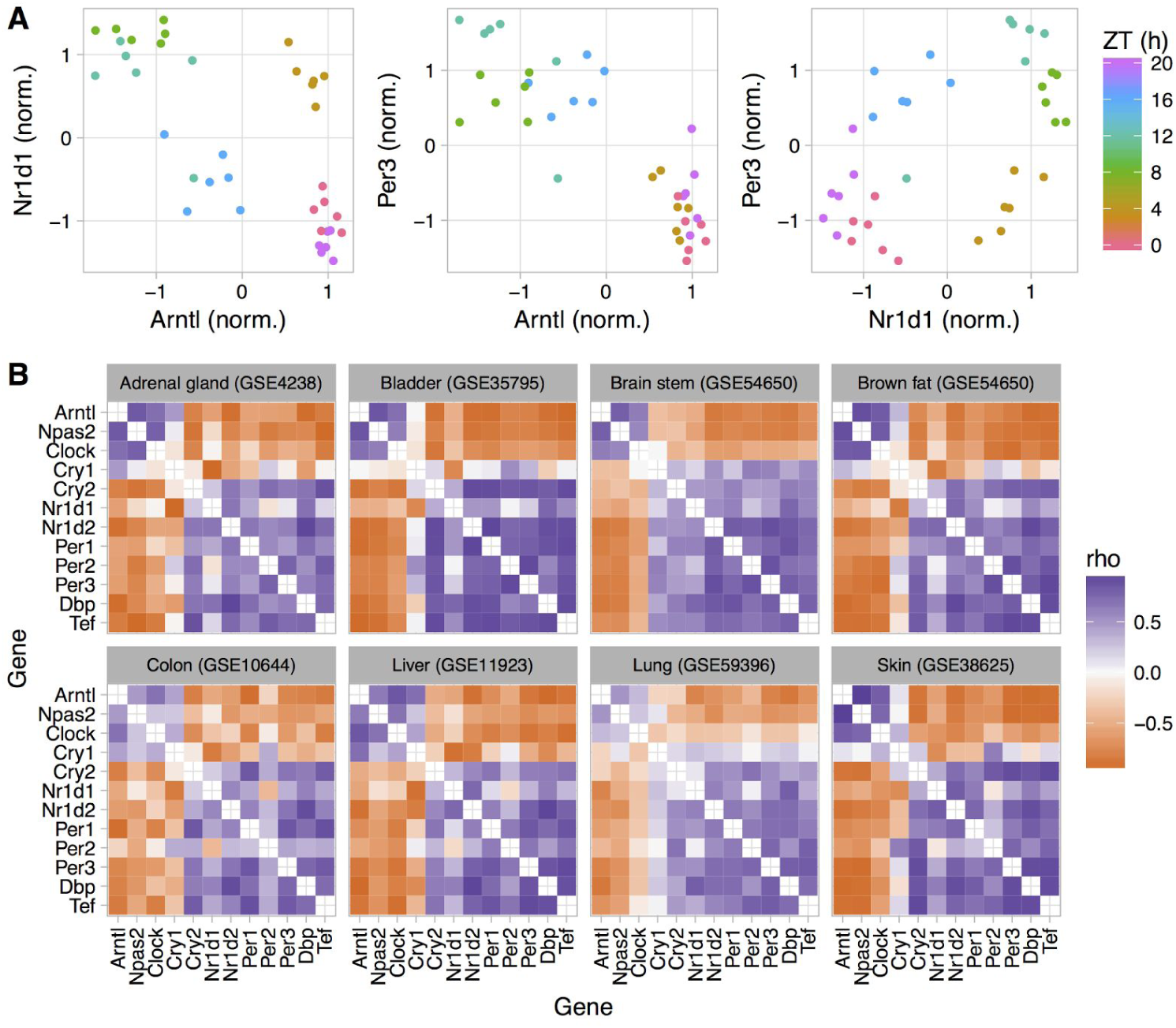
Consistent patterns of clock gene co-expression in various mouse organs. (**A**) Scatterplots of expression for three clock genes in lung (GSE59396). Each point is a sample, and the color indicates zeitgeber time, where ZT0 corresponds to “lights on.” Expression values of each gene were normalized to have mean zero and standard deviation one. (**B**) Heatmaps of Spearman correlation (“rho”) between each pair of clock genes in each dataset. Genes are ordered manually by a combination of name and known function in the clock. GSE54650 includes gene expression from 12 organs, however to maintain diversity in datasets, we used data from only two organs.

For each dataset, we calculated the Spearman correlation between expression values (over all samples) of each pair of genes. The pattern of correlations was highly similar across datasets and revealed two groups of genes, where the genes within a group tended to be positively correlated with each other and negatively correlated with genes in the other group (Fig. 1B). Genes in the first group (Arntl, Npas2, and Clock), which are known to form the positive arm of the clock *(41)*, peaked in expression shortly before zeitgeber time 0 (ZT0, which corresponds to time of lights on or sunrise; Fig. S1). Genes in the second group (Cry2, Nr1d1, Nr1d2, Per1, Per2, Per3, Dbp, and Tef), which are known to form the negative arms of the clock, peaked in expression near ZT10. Cry1, which appeared to be part of the first group in some datasets and the second group in others, tended to peak in expression around ZT18. These results indicate that the progression of the circadian clock in mice produces a consistent pattern of correlations in expression between clock genes. The pattern does not depend on the absolute phasing of clock gene expression relative to time of day. Consequently, the pattern is not affected by phase shifts, such as those caused by temporally restricted feeding *(42)* (Fig. S2).

Most computational methods for quantifying circadian rhythmicity and inferring the status of the clock require that samples be acquired over the entire 24-h cycle. Because our approach does not attempt to infer oscillations, we wondered if it would be robust to partial coverage of the 24-h cycle. We therefore analyzed clock gene co-expression in three of the previous datasets, in samples acquired during the first 8 h of the day (or subjective day) or the first 8 h of the night (or subjective night). In each dataset, the correlation pattern was preserved in both daytime and nighttime samples (Fig. S3). These results indicate that our approach can detect an active circadian clock in groups of samples without using time of day information, even if the samples’ coverage of the 24-h cycle is incomplete.

### Validation of the correlation pattern in humans

We next applied our approach to nine publicly available datasets of circadian transcriptome data from human tissues: one from skin *(43)*, two from brain *(44, 45)*, three from blood *(46–48)*, and three from cells cultured in vitro *(38, 49, 50)* (Table S1). The dataset from human skin consisted of samples taken at only three time-points for each subject (9:30am, 2:30pm, and 7:30pm). The datasets from human brain were based on postmortem tissue from multiple anatomical areas, and zeitgeber time for each sample was calculated using the respective donor’s date and time of death and geographic location. The datasets from human blood consisted of multiple samples taken throughout the 24-h cycle for each subject. The datasets from cells cultured in vitro were based on time-courses following synchronization by dexamethasone, serum, or alternating temperature cycles.

The patterns of clock gene co-expression in human tissues and cells were similar to the patterns in mice (Fig. 2 and Fig. S4), which is consistent with our previous findings of similar relative phasing of clock gene expression in mice and humans *(51)*. The pattern was less distinct in human blood (Fig. S5), likely because several clock genes show weak or no rhythmicity in expression in blood cells *(51)*. The strong pattern in human skin was due to clock gene co-expression both between the three time-points and between individuals at a given time-point (Fig. S6). Compared to co-expression patterns in mouse organs and human skin, those in human brain were somewhat weaker, which is consistent with the weaker circadian rhythmicity for clock genes in those two brain-specific datasets *(51)*.

**Figure 2.**
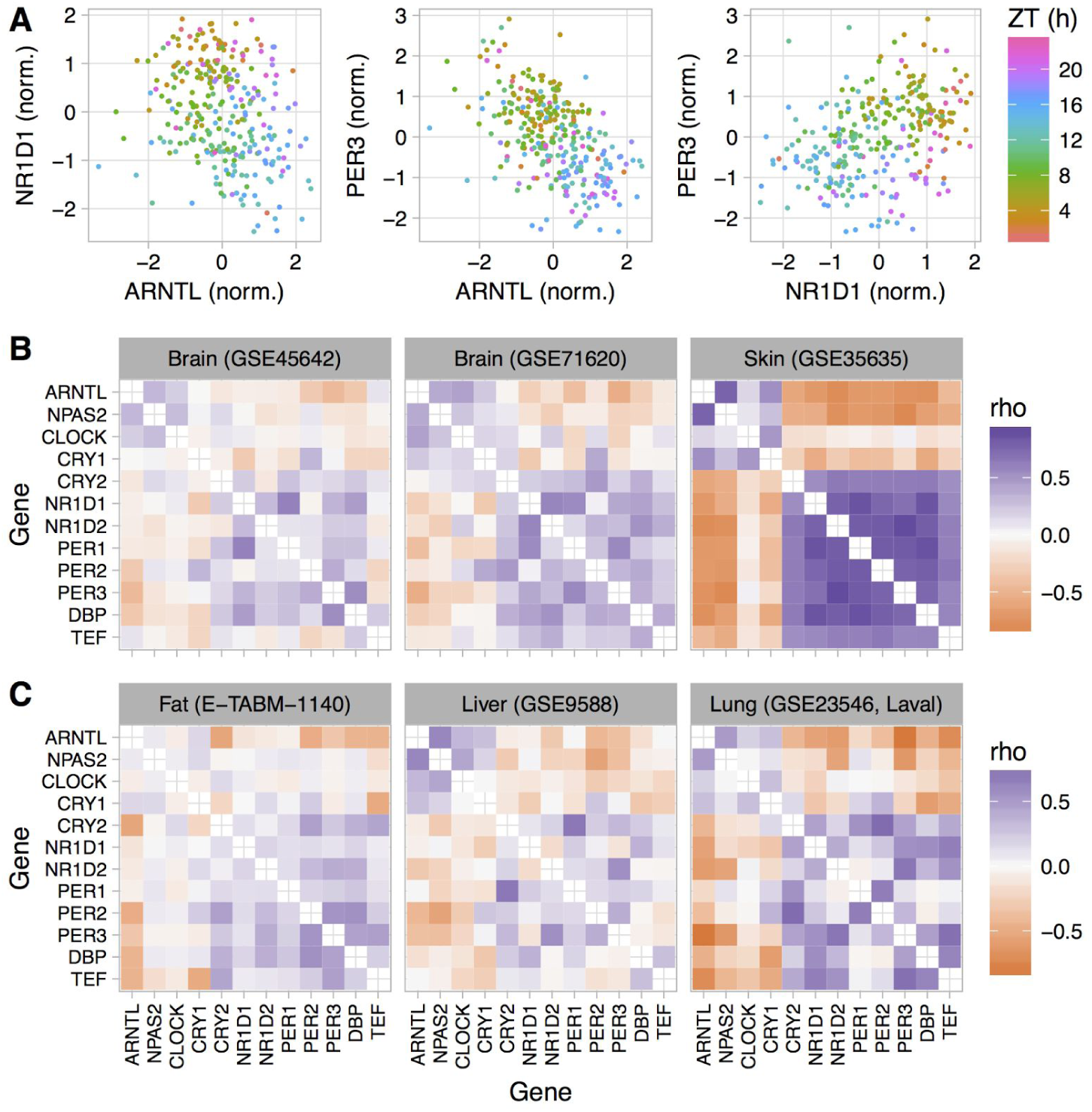
Consistent patterns of clock gene co-expression in healthy human organs, based on publicly available transcriptome data. (**A**) Scatterplots of expression for three clock genes in GSE71620 (gene expression measured in postmortem tissue, with zeitgeber time based on time of death). Each point is a sample, and the color corresponds to zeitgeber time, where ZT0 corresponds to sunrise. Expression values of each gene were normalized to have mean zero and standard deviation one. (**B**) Heatmaps of Spearman correlation between each pair of clock genes in three datasets designed to quantify circadian variation in gene expression. (**C**) Heatmaps for three datasets not designed to study circadian rhythms and in which samples are not labeled with time of day. Genes are in the same order as their mouse orthologs in Fig. 1.

To confirm our findings in a broader range of human organs, we analyzed five transcriptome datasets from healthy human lung, liver, skin, and adipose tissue *(52–56)* (Table S1). Samples from these datasets were not collected for the purpose of studying circadian rhythms and therefore are not labeled with time of day. Nonetheless, we observed the expected pattern of clock gene co-expression in each dataset (Fig. 2C and Fig. S7). We conclude that our approach can detect the signature of a functional circadian clock in a variety of human tissues in vitro and in vivo, even in datasets not designed to study circadian rhythms and with fewer than 100 samples.

### Aberrant patterns of clock gene co-expression in human cancer

To investigate patterns of clock gene co-expression in human cancer, we applied our approach to RNA-seq data collected by The Cancer Genome Atlas (TCGA) and reprocessed using the Rsubread package *(57)*. TCGA samples are from surgical resections performed prior to neoadjuvant treatment. The times of day of surgery are not available, but presumably most or all of the surgeries were performed during normal working hours. We analyzed data from the 12 cancer types that included at least 30 samples from adjacent non-tumor tissue (Table S1). For each cancer type, we calculated the Spearman correlations in expression between clock genes across all tumor samples and all non-tumor samples.

In non-tumor samples from most cancer types, we observed a similar pattern of clock gene co-expression as in the mouse and human circadian datasets (Fig. 3A-B and Fig. S8A). In contrast, in tumor samples from each cancer type, the pattern was weaker or absent. We observed the same trend when we restricted our analysis to only matched samples, i.e., samples from patients from whom both non-tumor and tumor samples were collected (Fig. S9). To confirm these findings, we analyzed eight additional datasets of gene expression in human cancer, four from liver and four from lung, each of which included matched tumor and adjacent non-tumor samples *(58–65)* (Table S1). As in the TCGA data, clock gene co-expression in tumor samples was perturbed relative to non-tumor samples (Fig. 3C and Fig. S8B).

**Figure 3.**
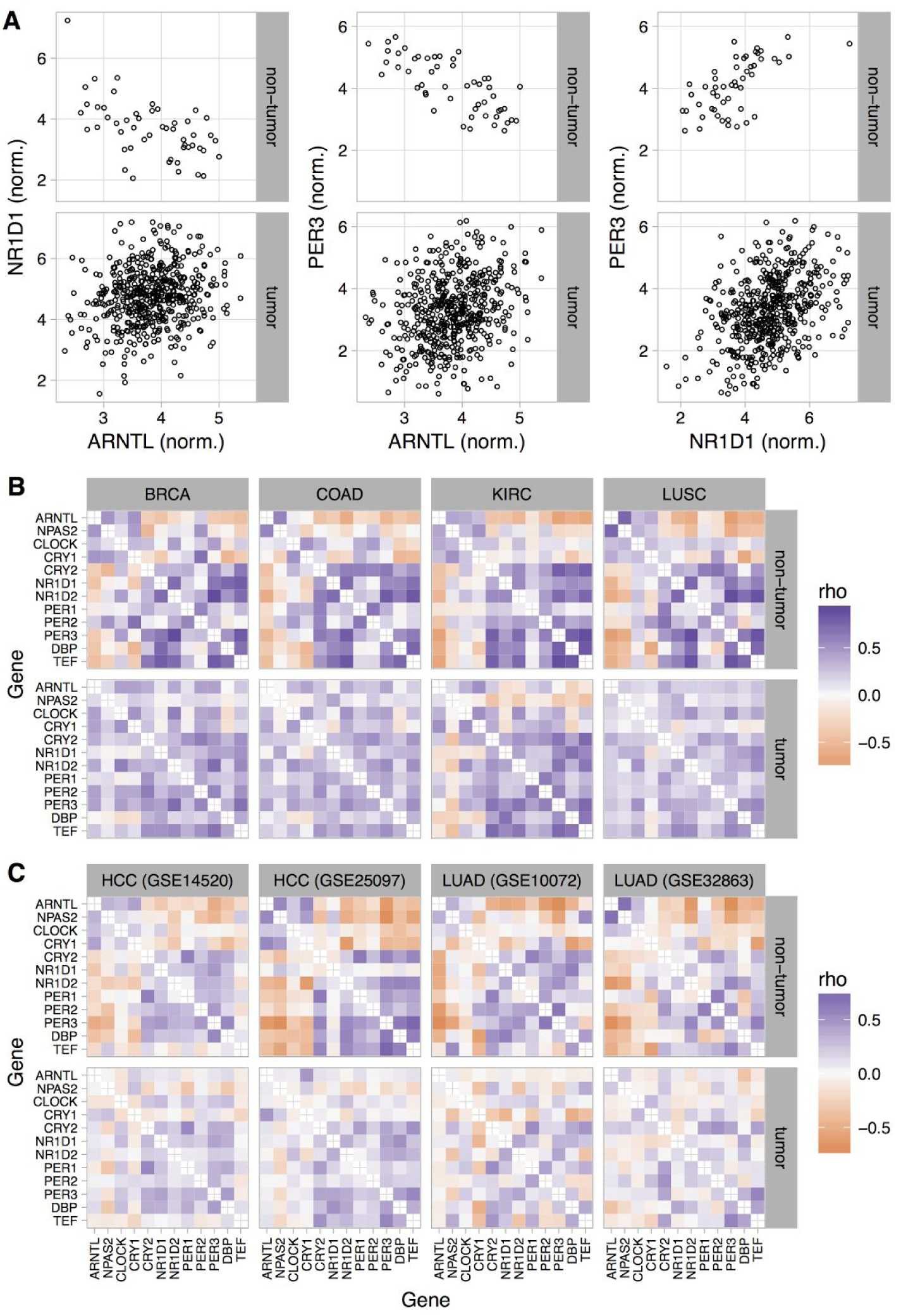
Loss of normal clock gene co-expression in tumor samples from various cancer types. (**A**) Pairwise scatterplots of expression for three clock genes in tumor and adjacent non-tumor samples from lung squamous cell carcinoma (LUSC) from TCGA. Each point is a sample. Expression values from RNA-seq data are shown in units of log2(tpm + 1). For ease of visualization, two tumor samples with very high normalized expression of ARNTL are not shown. (**B** and **C**) Heatmaps of Spearman correlation between each pair of clock genes in non-tumor and tumor samples from (**B**) four TCGA cancer types and (**C**) four additional datasets. TCGA cancer types shown are breast invasive cell carcinoma (BRCA), colon adenocarcinoma (COAD), kidney renal clear cell carcinoma (KIRC), and lung squamous cell carcinoma (LUSC). Additional datasets are from hepatocellular carcinoma (HCC) and lung adenocarcinoma (LUAD). In each dataset, all tumor samples are used, not only those with matched non-tumor samples.

To quantify the dysregulation of clock gene co-expression in human cancer, we first combined the eight mouse datasets in a fixed-effects meta-analysis (Fig. 4A and Methods) in order to construct a single “reference” correlation pattern (Fig. S10 and Table S2). For each of the 12 TCGA cancer types and each of the eight additional datasets of human cancer, we then calculated the Euclidean distances between the reference pattern and the non-tumor pattern and between the reference pattern and the tumor pattern. We refer to each of these distances as a clock correlation distance (CCD), and we refer to the difference between the tumor CCD and the non-tumor CCD as the delta clock correlation distance (ΔCCD). A positive ΔCCD indicates that the correlation pattern of the non-tumor samples is more similar to the reference than is the correlation pattern of the tumor samples.

**Figure 4.**
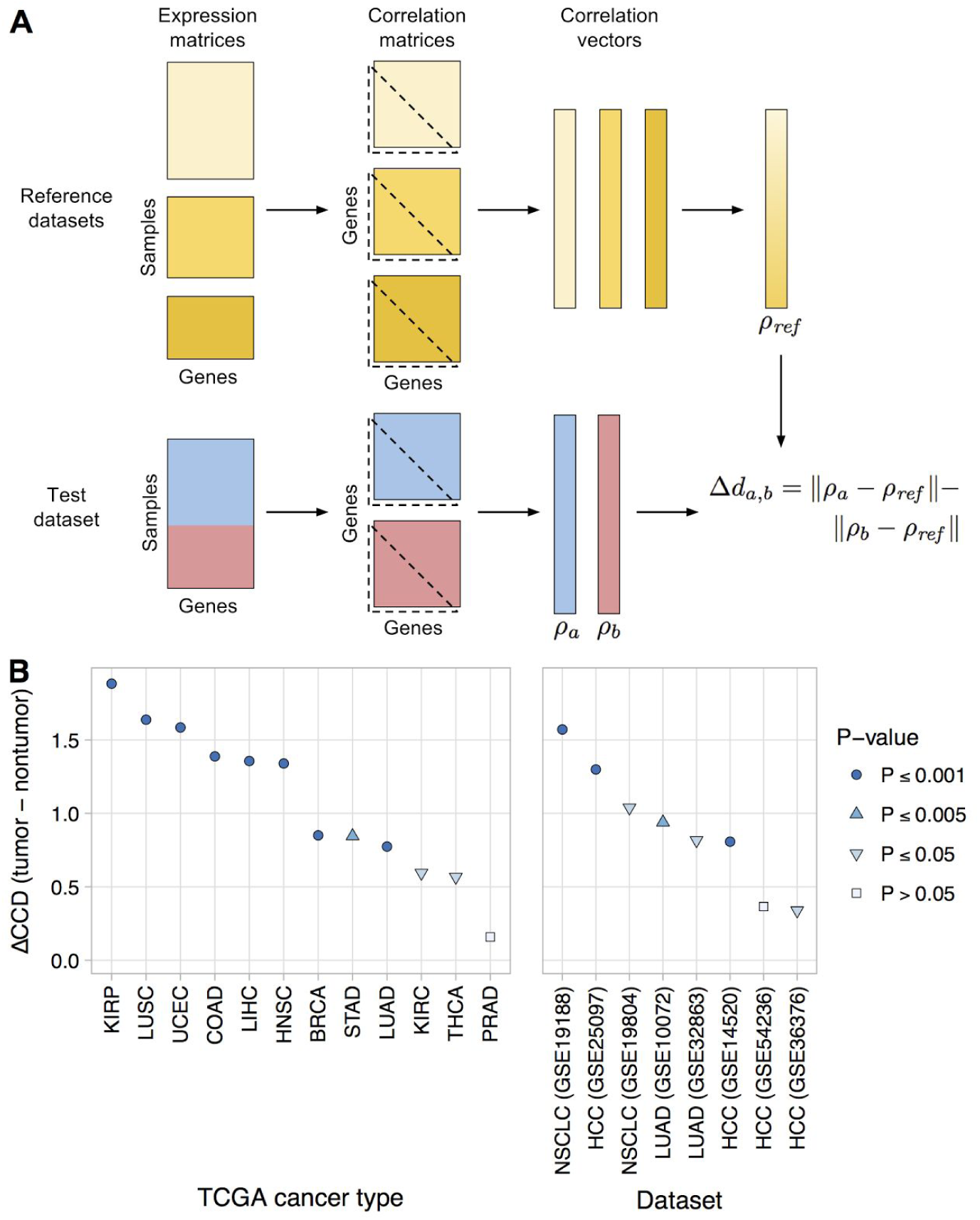
Quantifying dysregulation of clock gene co-expression in human cancer. (**A**) Schematic of procedure for comparing patterns of clock gene expression between two conditions in samples lacking time of day information. For details of procedure, see Methods. (**B**) Delta clock correlation distance (ΔCCD) between non-tumor and tumor samples in 12 TCGA cancer types and eight additional datasets. Additional datasets are from hepatocellular carcinoma (HCC), lung adenocarcinoma (LUAD), and non-small cell lung cancer (NSCLC). Positive ΔCCD indicates that the correlation pattern of the non-tumor samples is more similar to the mouse reference than is the correlation pattern of the tumor samples. P-values are one-sided and are based on 1000 permutations between the sample labels (non-tumor or tumor) and the gene expression values.

Consistent with the visualizations of clock gene co-expression, every TCGA cancer type and additional cancer dataset had a positive ΔCCD (Fig. 4B), as did the individual tumor grades in the TCGA data (Fig. S11). Among the three TCGA cancer types with the lowest ΔCCD, prostate adenocarcinoma had a relatively high non-tumor CCD (suggesting dysregulated clock gene co-expression even in non-tumor samples), whereas renal clear cell carcinoma and thyroid carcinoma each had a relatively low tumor CCD (Fig. S12). To evaluate the statistical significance of the ΔCCD, we permuted the sample labels (non-tumor or tumor) in each dataset and re-calculated the ΔCCD 1000 times. Based on this permutation testing, the observed ΔCCD for 11 of the 18 datasets had a one-sided P ≤ 0.001 (Fig. 4B). Overall, these results suggest that the circadian clock is dysregulated in a wide range of human cancers.

Tumors are a complex mixture of cancer cells and various non-cancerous cell types. The proportion of cancer cells in a tumor sample is called the tumor purity and is an important factor to consider in genomic analyses of bulk tumors *(66)*. We therefore examined the relationship between ΔCCD and tumor purity in the TCGA data. With the exception of thyroid carcinoma and prostate adenocarcinoma, ΔCCD and median tumor purity across TCGA cancer types were positively correlated (Fig. S13; Spearman correlation = 0.67, P = 0.059 by exact test). These findings suggest that at least in some cancer types, dysregulation of the circadian clock is stronger in cancer cells than in non-cancerous cells.

### Distinct patterns of clock gene expression in human cancer and mouse clock knockouts

We next investigated whether the clock dysregulation in human cancer resembled that caused by genetic mutations to core clock genes. We assembled seven datasets of circadian gene expression that included samples from wild-type mice and from mice in which at least one core clock gene was knocked out, either in the entire animal or in a specific cell type *(8, 42, 67–71)* (Table S1). For each dataset, we calculated the correlations in expression between pairs of clock genes in wild-type and mutant samples and calculated the ΔCCD (Fig. S14 and Fig. S15).

The two datasets with the highest ΔCCD (>50% higher than any ΔCCD we observed in human cancer) were those in which the mutant mice lacked not one, but two components of the clock (Cry1 and Cry2 in GSE13093; Nr1d1 and Nr1d2 in GSE34018). The ΔCCDs for the other five mutants were similar to or somewhat lower than the ΔCCDs we observed in human cancer. Given the smaller sample sizes compared to the human cancer datasets, the ΔCCDs for four of the other five mutants were not significantly greater than zero (one-sided P > 0.05 by permutation test).

To further compare clock dysregulation in human cancer and clock knockouts, we calculated differential expression of the clock genes between non-tumor and tumor samples and between wild-type and mutant samples (Fig. 5A). Differential expression in the clock knockouts was largely consistent with current understanding of the core clock. For example, knockout of Arntl (Bmal1, the primary transcriptional activator) tended to cause reduced expression (irrespective of time of day) of Nr1d1, Nr1d2, Per1, Per2, Per3, Dbp, and Tef, and increased or unchanged expression of the other clock genes. In the double knockout of Cry1 and Cry2 (two negative regulators of the clock), this pattern was reversed. Interestingly, neither of these patterns of differential expression was apparent in human cancer.

**Figure 5.**
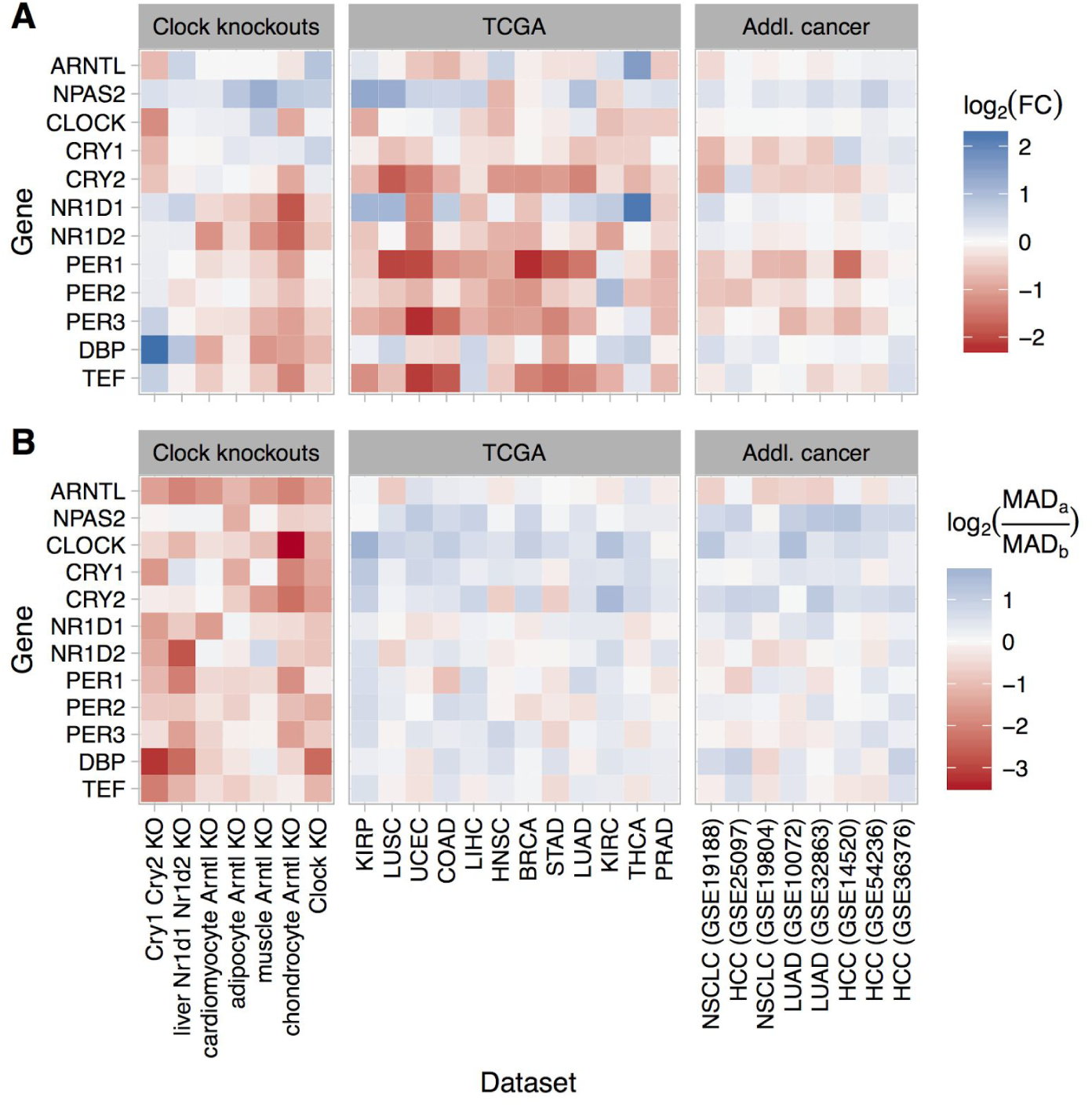
Dysregulation of the circadian clock in human cancer is distinct from that caused by knockout of the clock genes in mice. (**A**) Heatmaps of the estimated log2 fold-change in expression between tumor and non-tumor samples or between mutant and wild-type samples. A positive value indicates higher expression in tumor samples or mutant samples, respectively. (**B**) Heatmaps of the log2 ratio of the median absolute deviation (MAD) of expression in tumor compared to non-tumor samples and mutant compared to wild-type samples. In the legend, MAD_a_ refers to tumor or mutant samples, MAD_b_ refers to non-tumor or wild-type samples. A positive value indicates that the variation in expression of that gene is greater in tumor (or mutant) samples than in non-tumor (or wild-type) samples. For RNA-seq data (TCGA and chondrocyte Arntl KO), expression values were based on log2(tpm + 1). For microarray data, expression values were based on log-transformed, normalized intensity. Datasets are ordered by descending ΔCCD.

In the clock gene knockouts, rhythmic expression of the clock genes was reduced or lost (Fig. S16). Although it was not possible to directly quantify the rhythmicity of expression in the human cancer datasets, we reasoned that a proxy for rhythmicity could be the magnitude of variation in expression. Therefore, for each TCGA cancer type and each additional human cancer dataset, we calculated the median absolute deviation (MAD) in expression of the clock genes in non-tumor and tumor samples. We then compared the log_2_ ratios of MAD between tumor and non-tumor samples to the log_2_ ratios of MAD between mutant and wild-type samples from the clock gene knockout data (Fig. 5B). As expected, samples from clock gene knockouts showed widespread reductions in MAD compared to samples from wild-type mice. In contrast, human tumor samples tended to show similar or somewhat higher MAD compared to non-tumor samples. Taken together with the differential expression analysis, these results imply that clock dysregulation in human cancer is not due solely to loss of activity of one or more core clock genes.

### Dysregulation of broader circadian gene expression in human lung cancer

Finally, we explored how clock dysregulation in cancer might affect the circadian transcriptome. We obtained a set of 1,292 genes that were inferred by the CYCLOPS method to be rhythmic in a dataset from healthy human lung *(35)*. By applying WGCNA *(72)* to the same dataset, we grouped the 1,292 genes into five modules according to their co-expression (Fig. 6A). Based on DAVID *(73)* and consistent with the analysis of Anafi et al. *(35)*, the five modules were enriched for genes involved in various biological processes, including angiogenesis, phosphoprotein signaling, and alternative splicing (Table S3). We then used NetRep *(74)* to determine the extent to which each module was preserved in non-tumor and tumor samples from five datasets of human lung cancer (two from TCGA and three from NCBI GEO). Notably, all five datasets had ≤65 non-tumor samples and the datasets from NCBI GEO had ≤91 tumor samples, too few for CYCLOPS to produce a reliable ordering. Based on this analysis, four of five modules (comprising 1,256 of 1,292 rhythmic genes) were significantly more strongly preserved in non-tumor compared to tumor samples (Fig. 6B, P ≤ 0.001 for at least two of seven preservation statistics in at least four of five datasets).

**Figure 6.**
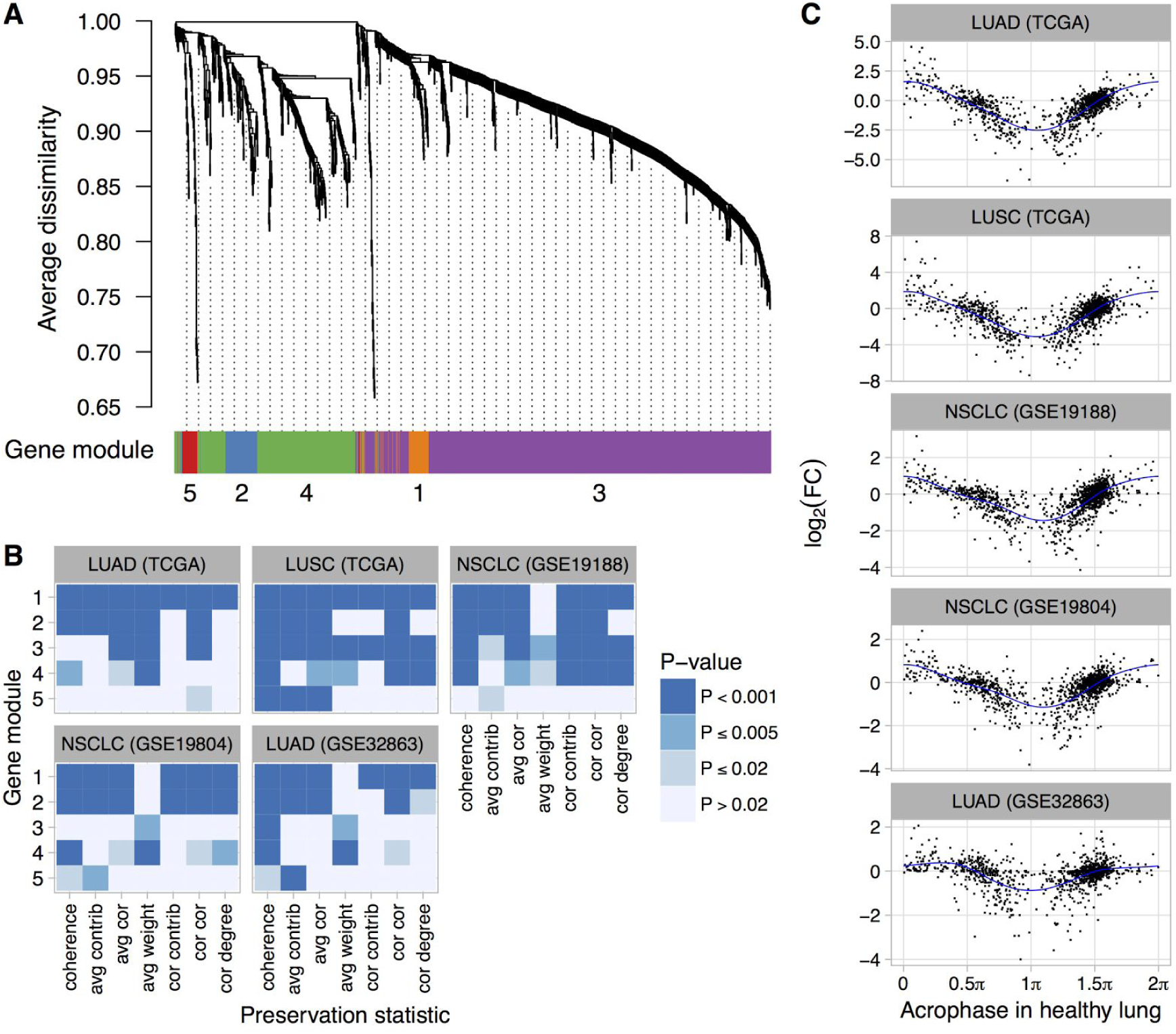
Perturbed expression of normally rhythmic genes in human lung cancer. (**A**) Hierarchical clustering of 1,292 genes inferred to have a circadian rhythm in healthy human lung. Gene modules were defined by following the procedure recommended by WGCNA. The number of genes in modules 1-5 are 62, 75, 828, 291, and 36. (**B**) Statistical significance of differential module preservation between non-tumor and tumor samples for seven preservation statistics for each module in each dataset. One-sided p-values are based on 1000 permutations of the sample labels in the respective test dataset. (**C**) Scatterplots of log_2_ fold-change in each lung cancer dataset vs. the phase of peak expression (acrophase, in radians) in healthy lung. Acrophase π is defined to be the circular mean of the acrophases of the circadian clock-driven PAR bZip transcription factors DBP, TEF, and HLF. Each point corresponds to one of the 1,292 genes. Each blue curve indicates a fit to a periodic smoothing spline. The circular mean of the trough of the spline fits is 1.06 π.

We also quantified differential expression of the 1,292 normally rhythmic genes between non-tumor and tumor samples in the five lung cancer datasets, and examined the relationship between each gene’s log_2_ fold-change and its phase of peak expression (acrophase) in healthy lung as inferred by CYCLOPS. Remarkably, we observed a consistent pattern, in which genes with an acrophase near π tended to show the strongest reductions in expression in tumor samples (Fig. 6C). Using Phase Set Enrichment Analysis *(75)*, we found that these genes were particularly enriched for involvement in G protein-coupled receptor signaling (Table S4). Taken together, these findings suggest that dysregulation of the circadian clock in human lung cancer is accompanied by systematic changes in broader circadian gene expression.

## Discussion

Increasing evidence has suggested that systemic disruption of the circadian clock can promote tumor development and that components of a tumor can disrupt the circadian clock. Until now, however, whether the clock is functional in primary human tumors has been unclear. Here we developed a simple method to probe clock function based on the co-expression of a small set of clock genes. By applying the method to publicly available cancer transcriptome data, we uncovered dysregulation of the circadian clock in multiple types of human cancer. Furthermore, our analysis of human lung cancer suggests that dysregulation of the clock is accompanied by large-scale changes in circadian gene expression.

Our approach for detecting a functional circadian clock relies on three principles. First, we use prior knowledge of clock genes and clock-controlled genes. Second, we account for the fact that the clock is defined not by a static condition, but by a dynamic cycle. Our approach thus exploits the co-expression of clock genes that arises from (1) different genes having rhythms with different circadian phases and (2) different samples being taken from different points in the cycle. Finally, our method does not attempt to infer an oscillatory pattern, but instead uses only the statistical correlations in expression between pairs of genes. The underlying assumption of the ΔCCD is that perturbations to the clock will alter the relative phases and/or signal-to-noise ratios of rhythms in clock gene expression and thereby alter the correlations. Although the correlation matrix only captures part of the complex relationship between genes, it is intuitive, simple to calculate, and requires relatively few samples (our results indicate that 30 in humans may be sufficient). Altogether, these principles enable our method to detect dysregulation of the clock in groups of samples whose times of day of acquisition are unknown and whose coverage of the 24-h cycle is incomplete. Consequently, the ΔCCD should be a valuable complement to methods designed to infer rhythms in omics data, such as CYCLOPS *(35)*.

Despite these advantages, our method does have limitations. First, a large ΔCCD does not exclude the possibility that the clock genes are still rhythmic, but instead implies that if rhythms are present, the phase relationships between genes are greatly altered relative to a normally functioning clock. Similarly, despite altered expression and co-expression levels, some of the genes rhythmic in healthy lung may still be rhythmic in lung cancer. Second, the ΔCCD is insensitive to the alignment of the circadian clock to time of day, and so cannot detect a phase difference between conditions, such as that recently observed during manic episodes of bipolar disorder *(76)*. This limitation, however, allowed us to readily compare clock gene co-expression between mice and humans, despite the circadian phase difference between the two species *(51)*. Third, the ΔCCD is invariant to the relative levels of gene expression between conditions, which is why we complemented our analysis of clock gene co-expression with analysis of differential expression and differential variability. Fourth, transcription is only one facet of the core clock mechanism, and perturbations to post-translational modification or degradation of clock proteins (if unaccompanied by changes in clock gene expression) would not be detected by our approach. Finally, because the ΔCCD relies on co-expression across samples, it does not immediately lend itself to quantifying clock disruption in single samples. In the future, it may be possible to complement the ΔCCD and assess clock function in some datasets by directly comparing matched samples from the same patient.

In healthy tissues in vivo, the circadian clocks of individual cells are entrained and oscillating together, which allows bulk measurements to contain robust circadian signals. Consequently, the loss of a circadian signature in human tumors could result from dysfunction in either entrainment, the oscillator, or both. Dysfunction in entrainment would imply that the clocks in at least some of the cancer cells are out of sync with each other and therefore free running, i.e. ignoring zeitgeber signals. Dysfunction in the oscillator would imply that the clocks in at least some of the cancer cells are no longer “ticking” normally. Given the current data, which can be considered averaged clock gene expression from many cells, these scenarios cannot be distinguished. However, the moderate correlation between ΔCCD and tumor purity across cancer types leads us to speculate that the clocks in stromal and/or infiltrating immune cells may be operating normally. In the future, the nature and mechanisms of clock dysregulation in vivo may be unraveled through a combination of mathematical modeling *(77, 78)* and single-cell measurements. A separate matter not addressed here is how the cancer influences circadian rhythms in the rest of the body *(79)*, which may be relevant for optimizing the daily timing of anticancer treatments *(80)*.

Based on this study alone, which is observational, it is not possible to determine whether clock dysregulation is a cancer driver or passenger. Indeed, perturbations to co-expression in cancer are not limited to clock genes or to rhythmic genes, as previous work has found changes in co-expression and connectivity across the transcriptome *(81, 82)*. However, given the clock’s established role in regulating metabolism and a recent finding that stimulation of the clock inhibits tumor growth in melanoma *(26)*, our findings raise the possibility that clock dysregulation and manipulation of normal circadian physiology may be a cancer driver in multiple solid tissues. On the other hand, a functional circadian clock seems to be required for growth of acute myeloid leukemia cells *(83)*, so further work is necessary to clarify this issue.

## Conclusions

Our findings suggest that dysregulation of the circadian clock is a common means by which human cancers achieve unrestrained growth and division. Thus, restoring clock function could be a viable therapeutic strategy in a wide range of cancer types. Given recent work on the effects of natural light exposure *(84)* and time-restricted feeding *(85, 86)*, such a therapeutic would not necessarily have to be pharmacological. In addition, given the practical challenges of studying circadian rhythms at the cellular level in humans, our method offers the possibility to quantify clock function in a wide range of human phenotypes using publicly available transcriptome data.

## Methods

### Selecting the datasets

We selected the datasets of circadian gene expression in mice (both for defining the reference pattern and for comparing clock gene knockouts to wild-type) to represent multiple organs, light-dark regimens, and microarray platforms. For circadian gene expression in humans, we included three datasets from blood, two from brain, and one from skin. The samples from blood and skin were obtained from living volunteers, whereas the samples from brain were obtained from postmortem donors who had died rapidly. For GSE45642 (human brain), we only included samples from control subjects (i.e., we excluded subjects with major depressive disorder). Zeitgeber times for samples from GSE56931 (human blood) were calculated as described previously *(87)*. For the TCGA data, we analyzed all cancer types that had at least 30 non-tumor samples (all of which also had at least 291 tumor samples). When analyzing clock gene expression in human cancer, unless otherwise noted, we included all tumor and non-tumor samples, not just those from patients from whom both non-tumor and tumor samples were collected. For details of the datasets, all of which are publicly available, see Table S1.

### Processing the gene expression data

For TCGA samples, we obtained the processed RNA-seq data (in units of transcripts per million, TPM, on a gene-level basis) and the corresponding metadata (cancer type, patient ID, etc.) from GSE62944 *(57)*. For E-MTAB-3428, we downloaded the RNA-seq read files from the European Nucleotide Archive, used Salmon to quantify transcript-level abundances in units of TPM *(88)*, then used the mapping between Ensembl Transcript IDs and Entrez Gene IDs to calculate gene-level abundances.

For the remaining datasets, raw (in the case of Affymetrix) or processed microarray data were obtained from NCBI GEO and processed using MetaPredict, which maps probes to Entrez Gene IDs and performs intra-study normalization and log-transformation *(89)*. MetaPredict processes raw Affymetrix data using RMA and customCDFs *(90, 91)*. As in our previous study, we used ComBat to reduce batch effects between anatomical areas in human brain and between subjects in human blood *(51, 92)*.

### Analyzing clock gene expression and co-expression

We first focused on the expression of 12 genes that are considered part of the core clock or are directly controlled by the clock and that show strong, consistently phased rhythms in a wide range of mouse organs *(3, 32)*. We calculated times of peak expression and strengths of circadian rhythmicity of expression in wild-type and mutant mice using ZeitZeiger *(32)*, with three knots for the periodic smoothing splines *(93)*.

We quantified the relationship between expression values of pairs of genes using the Spearman correlation (Spearman’s rho), which is rank-based and therefore invariant to monotonic transformations such as the logarithm and less sensitive to outliers than the Pearson correlation. Using the biweight midcorrelation, which is also robust to outliers, gave very similar results. All heatmaps of gene-gene correlations in this paper have the same mapping of correlation value to color, so they are directly visually comparable.

We calculated the reference Spearman correlation for each pair of genes (Table S2) using a fixed-effects meta-analysis *(94)* of the eight mouse datasets shown in Fig. 1. First, we applied the Fisher z-transformation (*arctanh*) to the correlations from each dataset. Then we calculated a weighted average of the transformed correlations, where the weight for dataset *i* was *n_i_* – 3 (corresponding to the inverse variance of the transformed correlation), where *n_i_* is the number of samples in dataset *i*. Finally, we applied the inverse transformation (*tanh*) to the weighted average.

To quantify the similarity in clock gene expression between two groups of samples (e.g., between the mouse reference and human tumor samples), we calculated the Euclidean distance between the respective Spearman correlation vectors, which contains all values in the strictly lower (or strictly upper) triangular part of the correlation matrix. Given a reference and a dataset with samples from two conditions, we calculated the Euclidean distances between the reference and each condition, which we call the clock correlation distances (CCDs). We then calculated the difference between these two distances, which indicates how much more similar to the reference one condition is than the other and which we refer to as the delta clock correlation distance (ΔCCD). Although here we used Euclidean distance, other distance metrics could be used as well.

To evaluate the statistical significance of the ΔCCD for a given dataset, we conducted the permutation test as follows: First, we permuted the relationship between the sample labels (e.g., non-tumor or tumor) and the gene expression values and recalculated the ΔCCD 1000 times, always keeping the reference fixed. We then calculated the one-sided p-value based on the number of permutations that gave a ΔCCD greater than or equal to the observed ΔCCD, adjusted using the method of Phipson and Smyth in the statmod R package *(95)*. Since we used the one-sided p-value, the alternative hypothesis was that non-tumor (or wild-type) is more similar to the reference than is tumor (or mutant).

To calculate the ΔCCD for individual tumor grades, we used the clinical metadata provided in GSE62944. We analyzed all combinations of TGCA cancer type and tumor grade that included at least 50 tumor samples. In each case, we calculated the ΔCCD using all non-tumor samples of the respective cancer type.

To compare ΔCCD and tumor purity, we used published consensus purity estimates for TCGA tumor samples *(66)*. The estimates are based on DNA methylation, somatic copy number variation, and the expression of immune genes and stromal genes (none of which are clock genes).

We quantified differential expression between tumor and non-tumor samples and between mutant and wild-type samples using limma and voom *(96, 97)*. As these techniques as designed for transcriptome data, we calculated differential expression using all measured genes. To ensure a fair comparison between human and mouse data, we ignored time of day information in the mouse samples. We quantified the variation in expression of clock genes in each dataset and condition using the median absolute deviation (MAD), which is less sensitive to outliers than the standard deviation.

### Analyzing circadian gene expression in human lung cancer

We obtained the set of rhythmic transcripts (identified by microarray probe ID) inferred by CYCLOPS to be rhythmic in healthy human lung *(35)*. This set contains transcripts whose abundance was well described by a sinusoidal function of CYCLOPS phase in samples from both the Laval and GRNG sites of GSE23546 (Bonferroni-corrected P < 0.05 and peak/trough ratio > 2) and whose orthologs were rhythmic in mouse lung. We mapped the microarray probes to Entrez Gene IDs and calculated the acrophase for each gene as the circular mean of the provided acrophase for the corresponding probes. Phase values inferred by CYCLOPS are relative, so Anafi et al. adopted the convention of setting phase π (in radians) to the average acrophase of the PAR bZip transcription factors (DBP, HLF, and TEF).

Because some of the module preservation statistics calculated by NetRep use the expression matrix directly, we used ComBat *(92)* to merge the expression data from the Laval and Groningen sites of GSE23546. We then followed WGCNA’s recommended procedure for identifying gene modules *(72)*. Briefly, starting with the merged expression data, we calculated the Spearman correlation matrix for the 1,292 genes, then used the “signed” method with a soft thresholding power of 12 to calculate the adjacency matrix, then calculated the topological overlap dissimilarity matrix. We used hierarchical clustering of the dissimilarity matrix and adaptive branch pruning to define modules of genes, followed by a procedure to merge closely related modules. We then used the DAVID web application (version 6.8) to discover functional categories enriched in each module *(73)*.

We used NetRep *(74)* to calculate seven module preservation statistics between healthy human lung and the non-tumor and tumor samples from five datasets of human lung cancer (LUAD and LUSC from TCGA and GSE19188, GSE19804, and GSE32863 from NCBI GEO). We excluded GSE10072, because it included expression levels for only 876 of the 1,292 genes (the other five included at least 1,199 of the 1,292 genes). When calculating module preservation, NetRep automatically removed any genes that were not measured in both healthy human lung and the group of samples to which it was being compared. For non-tumor and tumor samples from each dataset, we calculated the Spearman correlation matrix and adjacency matrix using the same procedure as for healthy human lung.

To evaluate the statistical significance of the difference in each module preservation statistic between non-tumor and tumor samples, we performed permutation testing and calculated one-sided p-values similarly to the procedure for ΔCCD, permuting the sample labels in the test dataset (non-tumor or tumor) and recalculating module preservation 1000 times (note this is different from the standard way WGCNA and NetRep perform permutations, which is to shuffle the gene labels).

Periodic smoothing splines of log_2_ fold-change as a function of acrophase in healthy lung were fit using ZeitZeiger *(32)*. For Phase Set Enrichment Analysis *(75)*, we used the “canonical pathways” gene sets from MSigDB *(98)* and looked for gene sets with a q-value < 0.1 against a uniform distribution and vector-average value within 0.4 radians of either 0.07 π or 1.06 π (the mean phases of the peak and trough of the spline fits, respectively). No gene sets met the criteria for the former.

## Abbreviations

ΔCCD: delta clock correlation distance
MAD: median absolute deviation
TCGA: The Cancer Genome Atlas
ZT: zeitgeber time
BRCA: breast invasive cell carcinoma
COAD: colon adenocarcinoma
HNSC: head and neck squamous cell carcinoma
KIRC: kidney renal clear cell carcinoma
KIRP: kidney renal papillary cell carcinoma
LIHC: liver hepatocellular carcinoma
LUAD: lung adenocarcinoma
LUSC: lung squamous cell carcinoma
PRAD: prostate adenocarcinoma
STAD: stomach adenocarcinoma
THCA: thyroid carcinoma
UCEC: uterine corpus endometrial carcinoma
TPM: transcripts per million

## Declarations

### Supplementary Materials

Figures S1-S16.

Table S1. Details of each dataset used in this study.

Table S2. Reference Spearman correlations between clock genes.

Table S3. Functional categories significantly enriched in circadian gene modules in human lung according to DAVID (Benjamini-Hochberg P ≤ 0.05).

Table S4. Gene sets in human lung with uniform q-value < 0.1 and vector-average value within 0.4 radians of 1.06 π according to Phase Set Enrichment Analysis.

## Acknowledgments

We thank Dvir Aran and the VUMC Editor’s Club for helpful comments on the manuscript.

## Availability of data and materials

All data and code to reproduce this study are available at https://figshare.com/s/2eaf11e88642418f7e81. The original gene expression data and metadata for all datasets are available from NCBI GEO or Array Express. A web application to calculate ΔCCD for one’s own gene expression data is available at https://hugheylab.shinyapps.io/deltaccd.

## Author contributions

JS performed the analysis and reviewed drafts of the paper. GC conceived and designed the analysis and reviewed drafts of the paper. JJH conceived and designed the analysis, performed the analysis, wrote the paper, and reviewed drafts of the paper. All authors read and approved the final manuscript.

## References

1. C. Dibner, U. Schibler, U. Albrecht, The Mammalian Circadian Timing System: Organization and Coordination of Central and Peripheral Clocks, Annu. Rev. Physiol. 72, 517–549 (2010).

2. S.-H. Yoo, S. Yamazaki, P. L. Lowrey, K. Shimomura, C. H. Ko, E. D. Buhr, S. M. Siepka, H.-K. Hong, W. J. Oh, O. J. Yoo, M. Menaker, J. S. Takahashi, PERIOD2::LUCIFERASE real-time reporting of circadian dynamics reveals persistent circadian oscillations in mouse peripheral tissues, Proc. Natl. Acad. Sci. U. S. A. 101, 5339–5346 (2004).

3. R. Zhang, N. F. Lahens, H. I. Ballance, M. E. Hughes, J. B. Hogenesch, A circadian gene expression atlas in mammals: implications for biology and medicine, Proc. Natl. Acad. Sci. U. S. A. 111, 16219–16224 (2014).

4. F. Damiola, N. L. Minh, N. Preitner, B. Kornmann, F. Fleury-Olela, U. Schibler, Restricted feeding uncouples circadian oscillators in peripheral tissues from the central pacemaker in the suprachiasmatic nucleus, Genes Dev. 14, 2950–2961 (2000).

5. G. Asher, H. Reinke, M. Altmeyer, M. Gutierrez-Arcelus, M. O. Hottiger, U. Schibler, Poly(ADP-ribose) polymerase 1 participates in the phase entrainment of circadian clocks to feeding, Cell 142, 943–953 (2010).

6. K. L. Eckel-Mahan, V. R. Patel, S. de Mateo, R. Orozco-Solis, N. J. Ceglia, S. Sahar, S. A. Dilag-Penilla, K. A. Dyar, P. Baldi, P. Sassone-Corsi, Reprogramming of the circadian clock by nutritional challenge, Cell 155, 1464–1478 (2013).

7. Y. Nakahata, S. Sahar, G. Astarita, M. Kaluzova, P. Sassone-Corsi, Circadian control of the NAD+ salvage pathway by CLOCK-SIRT1, Science 324, 654–657 (2009).

8. H. Cho, X. Zhao, M. Hatori, R. T. Yu, G. D. Barish, M. T. Lam, L.-W. Chong, L. DiTacchio, A. R. Atkins, C. K. Glass, C. Liddle, J. Auwerx, M. Downes, S. Panda, R. M. Evans, Regulation of circadian behaviour and metabolism by REV-ERB-α and REV-ERB-β, Nature 485, 123–127 (2012).

9. A. Neufeld-Cohen, M. S. Robles, R. Aviram, G. Manella, Y. Adamovich, B. Ladeuix, D. Nir, L. Rousso-Noori, Y. Kuperman, M. Golik, M. Mann, G. Asher, Circadian control of oscillations in mitochondrial rate-limiting enzymes and nutrient utilization by PERIOD proteins, Proc. Natl. Acad. Sci. U. S. A. 113, E1673–82 (2016).

10. T. Matsuo, S. Yamaguchi, S. Mitsui, A. Emi, F. Shimoda, H. Okamura, Control mechanism of the circadian clock for timing of cell division in vivo, Science 302, 255–259 (2003).

11. A. Gréchez-Cassiau, B. Rayet, F. Guillaumond, M. Teboul, F. Delaunay, The circadian clock component BMAL1 is a critical regulator of p21WAF1/CIP1 expression and hepatocyte proliferation, J. Biol. Chem. 283, 4535–4542 (2008).

12. M. Geyfman, V. Kumar, Q. Liu, R. Ruiz, W. Gordon, F. Espitia, E. Cam, S. E. Millar, P. Smyth, A. Ihler, J. S. Takahashi, B. Andersen, Brain and muscle Arnt-like protein-1 (BMAL1) controls circadian cell proliferation and susceptibility to UVB-induced DNA damage in the epidermis, Proc. Natl. Acad. Sci. U. S. A. 109, 11758–11763 (2012).

13. J. Bieler, R. Cannavo, K. Gustafson, C. Gobet, D. Gatfield, F. Naef, Robust synchronization of coupled circadian and cell cycle oscillators in single mammalian cells, Mol. Syst. Biol. 10, 739 (2014).

14. C. Feillet, P. Krusche, F. Tamanini, R. C. Janssens, M. J. Downey, P. Martin, M. Teboul, S. Saito, F. A. Lévi, T. Bretschneider, G. T. J. van der Horst, F. Delaunay, D. A. Rand, Phase locking and multiple oscillating attractors for the coupled mammalian clock and cell cycle, Proc. Natl. Acad. Sci. U. S. A. 111, 9828–9833 (2014).

15. T. Matsu-Ura, A. Dovzhenok, E. Aihara, J. Rood, H. Le, Y. Ren, A. E. Rosselot, T. Zhang, C. Lee, K. Obrietan, M. H. Montrose, S. Lim, S. R. Moore, C. I. Hong, Intercellular Coupling of the Cell Cycle and Circadian Clock in Adult Stem Cell Culture, Mol. Cell 64, 900–912 (2016).

16. E. S. Schernhammer, F. Laden, F. E. Speizer, W. C. Willett, D. J. Hunter, I. Kawachi, G. A. Colditz, Rotating night shifts and risk of breast cancer in women participating in the nurses health study, J. Natl. Cancer Inst. 93, 1563–1568 (2001).

17. E. S. Schernhammer, F. Laden, F. E. Speizer, W. C. Willett, D. J. Hunter, I. Kawachi, C. S. Fuchs, G. A. Colditz, Night-shift work and risk of colorectal cancer in the nurses’ health study, J. Natl. Cancer Inst. 95, 825–828 (2003).

18. E. S. Schernhammer, D. Feskanich, G. Liang, J. Han, Rotating night-shift work and lung cancer risk among female nurses in the United States, Am. J. Epidemiol. 178, 1434–1441 (2013).

19. L. R. Wegrzyn, R. M. Tamimi, B. A. Rosner, S. B. Brown, R. G. Stevens, A. H. Eliassen, F. Laden, W. C. Willett, S. E. Hankinson, E. S. Schernhammer, ROTATING NIGHT SHIFT WORK AND RISK OF BREAST CANCER IN THE NURSES’ HEALTH STUDIES, Am. J. Epidemiol. (2017), doi: 10.1093/aje/kwx140.

20. K. C. G. Van Dycke, W. Rodenburg, C. T. M. van Oostrom, L. W. M. van Kerkhof, J. L. A. Pennings, T. Roenneberg, H. van Steeg, G. T. J. van der Horst, Chronically Alternating Light Cycles Increase Breast Cancer Risk in Mice, Curr. Biol. 25, 1932–1937 (2015).

21. N. M. Kettner, H. Voicu, M. J. Finegold, C. Coarfa, A. Sreekumar, N. Putluri, C. A. Katchy, C. Lee, D. D. Moore, L. Fu, Circadian Homeostasis of Liver Metabolism Suppresses Hepatocarcinogenesis, Cancer Cell (2016), doi: 10.1016/j.ccell.2016.10.007.

22. T. Papagiannakopoulos, M. R. Bauer, S. M. Davidson, M. Heimann, L. Subbaraj, A. Bhutkar, J. Bartlebaugh, M. G. Vander Heiden, T. Jacks, Circadian Rhythm Disruption Promotes Lung Tumorigenesis, Cell Metab. 24, 324–331 (2016).

23. A. Relógio, P. Thomas, P. Medina-Pérez, S. Reischl, S. Bervoets, E. Gloc, P. Riemer, S. Mang-Fatehi, B. Maier, R. Schäfer, U. Leser, H. Herzel, A. Kramer, C. Sers, Ras-mediated deregulation of the circadian clock in cancer, PLoS Genet. 10, e1004338 (2014).

24. A. K. Michael, S. L. Harvey, P. J. Sammons, A. P. Anderson, H. M. Kopalle, A. H. Banham, C. L. Partch, Cancer/Testis Antigen PASD1 Silences the Circadian Clock, Mol. Cell 58, 743–754 (2015).

25. B. J. Altman, A. L. Hsieh, A. Sengupta, S. Y. Krishnanaiah, Z. E. Stine, Z. E. Walton, A. M. Gouw, A. Venkataraman, B. Li, P. Goraksha-Hicks, S. J. Diskin, D. I. Bellovin, M. C. Simon, J. C. Rathmell, M. A. Lazar, J. M. Maris, D. W. Felsher, J. B. Hogenesch, A. M. Weljie, C. V. Dang, MYC Disrupts the Circadian Clock and Metabolism in Cancer Cells, Cell Metab. 22, 1009–1019 (2015).

26. S. Kiessling, L. Beaulieu-Laroche, I. D. Blum, D. Landgraf, D. K. Welsh, K.-F. Storch, N. Labrecque, N. Cermakian, Enhancing circadian clock function in cancer cells inhibits tumor growth, BMC Biol. 15, 13 (2017).

27. A. Balsalobre, F. Damiola, U. Schibler, A serum shock induces circadian gene expression in mammalian tissue culture cells, Cell 93, 929–937 (1998).

28. S. Rossetti, J. Esposito, F. Corlazzoli, A. Gregorski, N. Sacchi, Entrainment of breast (cancer) epithelial cells detects distinct circadian oscillation patterns for clock and hormone receptor genes, Cell Cycle 11, 350–360 (2012).

29. S. Xiang, L. Mao, T. Duplessis, L. Yuan, R. Dauchy, E. Dauchy, D. E. Blask, T. Frasch, S. M. Hill, Oscillation of clock and clock controlled genes induced by serum shock in human breast epithelial and breast cancer cells: regulation by melatonin, Breast Cancer 6, 137–150 (2012).

30. G. Wu, R. C. Anafi, M. E. Hughes, K. Kornacker, J. B. Hogenesch, MetaCycle: an integrated R package to evaluate periodicity in large scale data, Bioinformatics 32, 3351–3353 (2016).

31. P. F. Thaben, P. O. Westermark, Differential rhythmicity: detecting altered rhythmicity in biological data, Bioinformatics 32, 2800–2808 (2016).

32. J. J. Hughey, T. Hastie, A. J. Butte, ZeitZeiger: supervised learning for high-dimensional data from an oscillatory system, Nucleic Acids Res. 44, e80 (2016).

33. C. Cadenas, L. van de Sandt, K. Edlund, M. Lohr, B. Hellwig, R. Marchan, M. Schmidt, J. Rahnenführer, H. Oster, J. G. Hengstler, Loss of circadian clock gene expression is associated with tumor progression in breast cancer, Cell Cycle 13, 3282–3291 (2014).

34. N. Leng, L.-F. Chu, C. Barry, Y. Li, J. Choi, X. Li, P. Jiang, R. M. Stewart, J. A. Thomson, C. Kendziorski, Oscope identifies oscillatory genes in unsynchronized single-cell RNA-seq experiments, Nat. Methods 12, 947–950 (2015).

35. R. C. Anafi, L. J. Francey, J. B. Hogenesch, J. Kim, CYCLOPS reveals human transcriptional rhythms in health and disease, Proc. Natl. Acad. Sci. U. S. A. (2017),doi: 10.1073/pnas.1619320114.

36. H. Oster, S. Damerow, R. A. Hut, G. Eichele, Transcriptional profiling in the adrenal gland reveals circadian regulation of hormone biosynthesis genes and nucleosome assembly genes, J. Biol. Rhythms 21, 350–361 (2006).

37. W. A. Hoogerwerf, M. Sinha, A. Conesa, B. A. Luxon, V. B. Shahinian, G. Cornélissen, F. Halberg, J. Bostwick, J. Timm, V. M. Cassone, Transcriptional profiling of mRNA expression in the mouse distal colon, Gastroenterology 135, 2019–2029 (2008).

38. M. E. Hughes, L. DiTacchio, K. R. Hayes, C. Vollmers, S. Pulivarthy, J. E. Baggs, S. Panda, J. B. Hogenesch, Harmonics of Circadian Gene Transcription in Mammals, PLoS Genet. 5, e1000442 (2009).

39. H. Negoro, A. Kanematsu, M. Doi, S. O. Suadicani, M. Matsuo, M. Imamura, T. Okinami, N. Nishikawa, T. Oura, S. Matsui, K. Seo, M. Tainaka, S. Urabe, E. Kiyokage, T. Todo, H. Okamura, Y. Tabata, O. Ogawa, Involvement of urinary bladder Connexin43 and the circadian clock in coordination of diurnal micturition rhythm, Nat. Commun. 3, 809 (2012).

40. J. A. Haspel, S. Chettimada, R. S. Shaik, J.-H. Chu, B. A. Raby, M. Cernadas, V. Carey, V. Process, G. M. Hunninghake, E. Ifedigbo, J. A. Lederer, J. Englert, A. Pelton, A. Coronata, L. E. Fredenburgh, A. M. K. Choi, Circadian rhythm reprogramming during lung inflammation, Nat. Commun. 5, 4753 (2014).

41. C. L. Partch, C. B. Green, J. S. Takahashi, Molecular architecture of the mammalian circadian clock, Trends Cell Biol. 24, 90–99 (2014).

42. C. Vollmers, S. Gill, L. DiTacchio, S. R. Pulivarthy, H. D. Le, S. Panda, Time of feeding and the intrinsic circadian clock drive rhythms in hepatic gene expression, Proc. Natl. Acad. Sci. U. S. A. 106, 21453–21458 (2009).

43. F. Spörl, S. Korge, K. Jürchott, M. Wunderskirchner, K. Schellenberg, S. Heins, A. Specht, C. Stoll, R. Klemz, B. Maier, H. Wenck, A. Schrader, D. Kunz, T. Blatt, A. Kramer, Krüppel-like factor 9 is a circadian transcription factor in human epidermis that controls proliferation of keratinocytes, Proc Natl Acad Sci USA 109, 10903–10908 (2012).

44. J. Z. Li, B. G. Bunney, F. Meng, M. H. Hagenauer, D. M. Walsh, M. P. Vawter, S. J. Evans, P. V. Choudary, P. Cartagena, J. D. Barchas, A. F. Schatzberg, E. G. Jones, R. M. Myers, S. J. Watson, H. Akil, W. E. Bunney, Circadian patterns of gene expression in the human brain and disruption in major depressive disorder, Proc Natl Acad Sci USA 110, 9950–9955 (2013).

45. C.-Y. Chen, R. W. Logan, T. Ma, D. A. Lewis, G. C. Tseng, E. Sibille, C. A. McClung, Effects of aging on circadian patterns of gene expression in the human prefrontal cortex, Proc Natl Acad Sci USA 113, 206–211 (2016).

46. C. S. Möller-Levet, S. N. Archer, G. Bucca, E. E. Laing, A. Slak, R. Kabiljo, J. C. Y. Lo, N. Santhi, M. von Schantz, C. P. Smith, D.-J. Dijk, Effects of insufficient sleep on circadian rhythmicity and expression amplitude of the human blood transcriptome, Proc Natl Acad Sci USA 110, E1132–E1141 (2013).

47. S. N. Archer, E. E. Laing, C. S. Möller-Levet, D. R. van der Veen, G. Bucca, A. S. Lazar, N. Santhi, A. Slak, R. Kabiljo, M. von Schantz, C. P. Smith, D.-J. Dijk, Mistimed sleep disrupts circadian regulation of the human transcriptome, Proc Natl Acad Sci USA 111, E682–E691 (2014).

48. E. S. Arnardottir, E. V. Nikonova, K. R. Shockley, A. A. Podtelezhnikov, R. C. Anafi, K. Q. Tanis, G. Maislin, D. J. Stone, J. J. Renger, C. J. Winrow, A. I. Pack, Blood-Gene Expression Reveals Reduced Circadian Rhythmicity in Individuals Resistant to Sleep Deprivation, Sleep 37, 589–600 (2014).

49. P. Janich, K. Toufighi, G. Solanas, N. M. Luis, S. Minkwitz, L. Serrano, B. Lehner, S. A. Benitah, Human epidermal stem cell function is regulated by circadian oscillations, Cell Stem Cell 13, 745–753 (2013).

50. J. Hoffmann, L. Symul, A. Shostak, T. Fischer, F. Naef, M. Brunner, Non-circadian expression masking clock-driven weak transcription rhythms in U2OS cells, PLoS One 9, e102238 (2014).

51. J. J. Hughey, A. J. Butte, Differential Phasing between Circadian Clocks in the Brain and Peripheral Organs in Humans, J. Biol. Rhythms 31, 588–597 (2016).

52. E. E. Schadt, C. Molony, E. Chudin, K. Hao, X. Yang, P. Y. Lum, A. Kasarskis, B. Zhang, S. Wang, C. Suver, J. Zhu, J. Millstein, S. Sieberts, J. Lamb, D. GuhaThakurta, J. Derry, J. D. Storey, I. Avila-Campillo, M. J. Kruger, J. M. Johnson, C. A. Rohl, A. van Nas, M. Mehrabian, T. A. Drake, A. J. Lusis, R. C. Smith, F. P. Guengerich, S. C. Strom, E. Schuetz, T. H. Rushmore, R. Ulrich, Mapping the genetic architecture of gene expression in human liver, PLoS Biol. 6, e107 (2008).

53. F. Innocenti, G. M. Cooper, I. B. Stanaway, E. R. Gamazon, J. D. Smith, S. Mirkov, J. Ramirez, W. Liu, Y. S. Lin, C. Moloney, S. F. Aldred, N. D. Trinklein, E. Schuetz, D. A. Nickerson, K. E. Thummel, M. J. Rieder, A. E. Rettie, M. J. Ratain, N. J. Cox, C. D. Brown, Identification, replication, and functional fine-mapping of expression quantitative trait loci in primary human liver tissue, PLoS Genet. 7, e1002078 (2011).

54. Y. Bossé, D. S. Postma, D. D. Sin, M. Lamontagne, C. Couture, N. Gaudreault, P. Joubert, V. Wong, M. Elliott, M. van den Berge, C. A. Brandsma, C. Tribouley, V. Malkov, J. A. Tsou, G. J. Opiteck, J. C. Hogg, A. J. Sandford, W. Timens, P. D. Paré, M. Laviolette, Molecular signature of smoking in human lung tissues, Cancer Res. 72, 3753–3763 (2012).

55. E. Grundberg, K. S. Small, Å. K. Hedman, A. C. Nica, A. Buil, S. Keildson, J. T. Bell, T.-P. Yang, E. Meduri, A. Barrett, J. Nisbett, M. Sekowska, A. Wilk, S.-Y. Shin, D. Glass, M. Travers, J. L. Min, S. Ring, K. Ho, G. Thorleifsson, A. Kong, U. Thorsteindottir, C. Ainali, A. S. Dimas, N. Hassanali, C. Ingle, D. Knowles, M. Krestyaninova, C. E. Lowe, P. Di Meglio, S. B. Montgomery, L. Parts, S. Potter, G. Surdulescu, L. Tsaprouni, S. Tsoka, V. Bataille, R. Durbin, F. O. Nestle, S. O’Rahilly, N. Soranzo, C. M. Lindgren, K. T. Zondervan, K. R. Ahmadi, E. E. Schadt, K. Stefansson, G. D. Smith, M. I. McCarthy, P. Deloukas, E. T. Dermitzakis, T. D. Spector, Multiple Tissue Human Expression Resource (MuTHER) Consortium, Mapping cis- and trans-regulatory effects across multiple tissues in twins, Nat. Genet. 44, 1084–1089 (2012).

56. M. J. Bonder, S. Kasela, M. Kals, R. Tamm, K. Lokk, I. Barragan, W. A. Buurman, P. Deelen, J.-W. Greve, M. Ivanov, S. S. Rensen, J. V. van Vliet-Ostaptchouk, M. G. Wolfs, J. Fu, M. H. Hofker, C. Wijmenga, A. Zhernakova, M. Ingelman-Sundberg, L. Franke, L. Milani, Genetic and epigenetic regulation of gene expression in fetal and adult human livers, BMC Genomics 15, 860 (2014).

57. M. Rahman, L. K. Jackson, W. E. Johnson, D. Y. Li, A. H. Bild, S. R. Piccolo, Alternative preprocessing of RNA-Sequencing data in The Cancer Genome Atlas leads to improved analysis results, Bioinformatics 31, 3666–3672 (2015).

58. M. T. Landi, T. Dracheva, M. Rotunno, J. D. Figueroa, H. Liu, A. Dasgupta, F. E. Mann, J. Fukuoka, M. Hames, A. W. Bergen, S. E. Murphy, P. Yang, A. C. Pesatori, D. Consonni, P. A. Bertazzi, S. Wacholder, J. H. Shih, N. E. Caporaso, J. Jen, Gene expression signature of cigarette smoking and its role in lung adenocarcinoma development and survival, PLoS One 3, e1651 (2008).

59. J. Hou, J. Aerts, B. den Hamer, W. van Ijcken, M. den Bakker, P. Riegman, C. van der Leest, P. van der Spek, J. A. Foekens, H. C. Hoogsteden, F. Grosveld, S. Philipsen, Gene expression-based classification of non-small cell lung carcinomas and survival prediction, PLoS One 5, e10312 (2010).

60. T.-P. Lu, M.-H. Tsai, J.-M. Lee, C.-P. Hsu, P.-C. Chen, C.-W. Lin, J.-Y. Shih, P.-C. Yang, C. K. Hsiao, L.-C. Lai, E. Y. Chuang, Identification of a novel biomarker, SEMA5A, for non-small cell lung carcinoma in nonsmoking women, Cancer Epidemiol. Biomarkers Prev. 19, 2590–2597 (2010).

61. S. Roessler, H.-L. Jia, A. Budhu, M. Forgues, Q.-H. Ye, J.-S. Lee, S. S. Thorgeirsson, Z. Sun, Z.-Y. Tang, L.-X. Qin, X. W. Wang, A unique metastasis gene signature enables prediction of tumor relapse in early-stage hepatocellular carcinoma patients, Cancer Res. 70, 10202–10212 (2010).

62. J. R. Lamb, C. Zhang, T. Xie, K. Wang, B. Zhang, K. Hao, E. Chudin, H. B. Fraser, J. Millstein, M. Ferguson, C. Suver, I. Ivanovska, M. Scott, U. Philippar, D. Bansal, Z. Zhang, J. Burchard, R. Smith, D. Greenawalt, M. Cleary, J. Derry, A. Loboda, J. Watters, R. T. P. Poon, S. T. Fan, C. Yeung, N. P. Y. Lee, J. Guinney, C. Molony, V. Emilsson, C. Buser-Doepner, J. Zhu, S. Friend, M. Mao, P. M. Shaw, H. Dai, J. M. Luk, E. E. Schadt, Predictive genes in adjacent normal tissue are preferentially altered by sCNV during tumorigenesis in liver cancer and may rate limiting, PLoS One 6, e20090 (2011).

63. S. A. Selamat, B. S. Chung, L. Girard, W. Zhang, Y. Zhang, M. Campan, K. D. Siegmund,M. N. Koss, J. A. Hagen, W. L. Lam, S. Lam, A. F. Gazdar, I. A. Laird-Offringa, Genome-scale analysis of DNA methylation in lung adenocarcinoma and integration with mRNA expression, Genome Res. 22, 1197–1211 (2012).

64. H.-Y. Lim, I. Sohn, S. Deng, J. Lee, S. H. Jung, M. Mao, J. Xu, K. Wang, S. Shi, J. W. Joh, Y. L. Choi, C.-K. Park, Prediction of disease-free survival in hepatocellular carcinoma by gene expression profiling, Ann. Surg. Oncol. 20, 3747–3753 (2013).

65. E. Villa, R. Critelli, B. Lei, G. Marzocchi, C. Cammà, G. Giannelli, P. Pontisso, G. Cabibbo, M. Enea, S. Colopi, C. Caporali, T. Pollicino, F. Milosa, A. Karampatou, P. Todesca, E. Bertolini, L. Maccio, M. L. Martinez-Chantar, E. Turola, M. Del Buono, N. De Maria, S. Ballestri, F. Schepis, P. Loria, G. Enrico Gerunda, L. Losi, U. Cillo, Neoangiogenesis-related genes are hallmarks of fast-growing hepatocellular carcinomas and worst survival. Results from a prospective study, Gut 65, 861–869 (2016).

66. D. Aran, M. Sirota, A. J. Butte, Systematic pan-cancer analysis of tumour purity, Nat.Commun. 6, 8971 (2015).

67. S. Nikolaeva, S. Pradervand, G. Centeno, V. Zavadova, N. Tokonami, M. Maillard, O. Bonny, D. Firsov, The circadian clock modulates renal sodium handling, J. Am. Soc. Nephrol. 23, 1019–1026 (2012).

68. G. K. Paschos, S. Ibrahim, W.-L. Song, T. Kunieda, G. Grant, T. M. Reyes, C. A. Bradfield, C. H. Vaughan, M. Eiden, M. Masoodi, J. L. Griffin, F. Wang, J. A. Lawson, G. A. Fitzgerald, Obesity in mice with adipocyte-specific deletion of clock component Arntl, Nat. Med. 18, 1768–1777 (2012).

69. K. A. Dyar, S. Ciciliot, L. E. Wright, R. S. Biensø, G. M. Tagliazucchi, V. R. Patel, M. Forcato, M. I. P. Paz, A. Gudiksen, F. Solagna, M. Albiero, I. Moretti, K. L. Eckel-Mahan, P. Baldi, P. Sassone-Corsi, R. Rizzuto, S. Bicciato, H. Pilegaard, B. Blaauw, S. Schiaffino, Muscle insulin sensitivity and glucose metabolism are controlled by the intrinsic muscle clock, Mol Metab 3, 29–41 (2014).

70. M. E. Young, R. A. Brewer, R. A. Peliciari-Garcia, H. E. Collins, L. He, T. L. Birky, B. W. Peden, E. G. Thompson, B.-J. Ammons, M. S. Bray, J. C. Chatham, A. R. Wende, Q. Yang,C.-W. Chow, T. A. Martino, K. L. Gamble, Cardiomyocyte-specific BMAL1 plays critical roles in metabolism, signaling, and maintenance of contractile function of the heart, J. Biol. Rhythms 29, 257–276 (2014).

71. M. Dudek, N. Gossan, N. Yang, H.-J. Im, J. P. D. Ruckshanthi, H. Yoshitane, X. Li, D. Jin, P. Wang, M. Boudiffa, I. Bellantuono, Y. Fukada, R. P. Boot-Handford, Q.-J. Meng, The chondrocyte clock gene Bmal1 controls cartilage homeostasis and integrity, J. Clin. Invest. 126, 365–376 (2016).

72. P. Langfelder, S. Horvath, WGCNA: an R package for weighted correlation network analysis, BMC Bioinformatics 9, 559 (2008).

73. D. W. Huang, B. T. Sherman, R. A. Lempicki, Systematic and integrative analysis of large gene lists using DAVID bioinformatics resources, Nat. Protoc. 4, 44–57 (2009).

74. S. C. Ritchie, S. Watts, L. G. Fearnley, K. E. Holt, G. Abraham, M. Inouye, A Scalable Permutation Approach Reveals Replication and Preservation Patterns of Network Modules in Large Datasets, Cell Syst 3, 71–82 (2016).

75. R. Zhang, A. A. Podtelezhnikov, J. B. Hogenesch, R. C. Anafi, Discovering Biology in Periodic Data through Phase Set Enrichment Analysis (PSEA), J. Biol. Rhythms 31, 244–257 (2016).

76. J.-H. Moon, C.-H. Cho, G. H. Son, D. Geum, S. Chung, H. Kim, S.-G. Kang, Y.-M. Park, H.-K. Yoon, L. Kim, H.-J. Jee, H. An, D. F. Kripke, H.-J. Lee, Advanced Circadian Phase in Mania and Delayed Circadian Phase in Mixed Mania and Depression Returned to Normal after Treatment of Bipolar Disorder, EBioMedicine 11, 285–295 (2016).

77. J. K. Kim, D. B. Forger, A mechanism for robust circadian timekeeping via stoichiometric balance, Mol. Syst. Biol. 8, 630 (2012).

78. S. Lück, K. Thurley, P. F. Thaben, P. O. Westermark, Rhythmic degradation explains and unifies circadian transcriptome and proteome data, Cell Rep. 9, 741–751 (2014).

79. S. Masri, T. Papagiannakopoulos, K. Kinouchi, Y. Liu, M. Cervantes, P. Baldi, T. Jacks, P. Sassone-Corsi, Lung Adenocarcinoma Distally Rewires Hepatic Circadian Homeostasis, Cell 165, 896–909 (2016).

80. S. Dulong, A. Ballesta, A. Okyar, F. Lévi, Identification of Circadian Determinants of Cancer Chronotherapy through In Vitro Chronopharmacology and Mathematical Modeling, Mol. Cancer Ther. 14, 2154–2164 (2015).

81. J. West, G. Bianconi, S. Severini, A. E. Teschendorff, Differential network entropy reveals cancer system hallmarks, Sci. Rep. 2, 802 (2012).

82. R. Anglani, T. M. Creanza, V. C. Liuzzi, A. Piepoli, A. Panza, A. Andriulli, N. Ancona, Loss of connectivity in cancer co-expression networks, PLoS One 9, e87075 (2014).

83. R. V. Puram, M. S. Kowalczyk, C. G. de Boer, R. K. Schneider, P. G. Miller, M. McConkey, Z. Tothova, H. Tejero, D. Heckl, M. Järås, M. C. Chen, H. Li, A. Tamayo, G. S. Cowley, O. Rozenblatt-Rosen, F. Al-Shahrour, A. Regev, B. L. Ebert, Core Circadian Clock Genes Regulate Leukemia Stem Cells in AML, Cell 165, 303–316 (2016).

84. E. R. Stothard, A. W. McHill, C. M. Depner, B. R. Birks, T. M. Moehlman, H. K. Ritchie, J. R. Guzzetti, E. D. Chinoy, M. K. LeBourgeois, J. Axelsson, K. P. Wright Jr, Circadian Entrainment to the Natural Light-Dark Cycle across Seasons and the Weekend, Curr. Biol. (2017), doi: 10.1016/j.cub.2016.12.041.

85. M. Hatori, C. Vollmers, A. Zarrinpar, L. DiTacchio, E. A. Bushong, S. Gill, M. Leblanc, A. Chaix, M. Joens, J. A. J. Fitzpatrick, M. H. Ellisman, S. Panda, Time-restricted feeding without reducing caloric intake prevents metabolic diseases in mice fed a high-fat diet, Cell Metab. 15, 848–860 (2012).

86. A. Chaix, A. Zarrinpar, P. Miu, S. Panda, Time-restricted feeding is a preventative and therapeutic intervention against diverse nutritional challenges, Cell Metab. 20, 991–1005 (2014).

87. J. J. Hughey, Machine learning identifies a compact gene set for monitoring the circadian clock in human blood, Genome Med. 9, 19 (2017).

88. R. Patro, G. Duggal, M. I. Love, R. A. Irizarry, C. Kingsford, Salmon provides fast and bias-aware quantification of transcript expression, Nat. Methods 14, 417–419 (2017).

89. J. J. Hughey, A. J. Butte, Robust meta-analysis of gene expression using the elastic net, Nucleic Acids Res. 43, e79 (2015).

90. R. A. Irizarry, B. Hobbs, F. Collin, Y. D. Beazer-Barclay, K. J. Antonellis, U. Scherf, T. P. Speed, Exploration, normalization, and summaries of high density oligonucleotide array probe level data, Biostatistics 4, 249–264 (2003).

91. M. Dai, P. Wang, A. D. Boyd, G. Kostov, B. Athey, E. G. Jones, W. E. Bunney, R. M. Myers, T. P. Speed, H. Akil, S. J. Watson, F. Meng, Evolving gene/transcript definitions significantly alter the interpretation of GeneChip data, Nucleic Acids Res. 33, e175 (2005).

92. W. E. Johnson, C. Li, A. Rabinovic, Adjusting batch effects in microarray expression data using empirical Bayes methods, Biostatistics 8, 118–127 (2007).

93. N. E. Helwig, P. Ma, Fast and stable multiple smoothing parameter selection in smoothing spline analysis of variance models with large samples, J. Comput. Graph. Stat. 24, 715–732 (2014).

94. L. V. Hedges, I. Olkin, Statistical Methods for Meta-Analysis (Academic Press, 1985).

95. B. Phipson, G. K. Smyth, Permutation P-values should never be zero: calculating exact P-values when permutations are randomly drawn, Stat. Appl. Genet. Mol. Biol. 9, Article39 (2010).

96. G. K. Smyth, Linear models and empirical bayes methods for assessing differential expression in microarray experiments, Stat. Appl. Genet. Mol. Biol. 3, Article3 (2004).

97. C. W. Law, Y. Chen, W. Shi, G. K. Smyth, voom: Precision weights unlock linear model analysis tools for RNA-seq read counts, Genome Biol. 15, R29 (2014).

98. A. Liberzon, A. Subramanian, R. Pinchback, H. Thorvaldsdóttir, P. Tamayo, J. P. Mesirov, Molecular signatures database (MSigDB) 3.0, Bioinformatics 27, 1739–1740 (2011).

